# New *Callicarpa* (Lamiaceae) taxa: Two species and a natural hybrid from Hahajima Island, Ogasawara Islands, Japan

**DOI:** 10.1101/2025.05.18.654768

**Authors:** Kyoko Sugai, Suzuki Setsuko, Kayo Hayama, Hidetoshi Kato

## Abstract

Two newly identified species of *Callicarpa* (Lamiaceae), *C. boninensis* and *C. hahajimensis*, along with a new natural hybrid, *C*. × *chibusensis* are described, all of which were identified on Hahajima Island, Ogasawara Islands, Japan. A comprehensive key to *Callicarpa* species in the Ogasawara Islands is also provided. *Callicarpa boninensis*, morphologically most similar to *C. subpubescens*, is distinguished by persistent stellate hairs on mature leaves and elongated petioles. *Callicarpa hahajimensis*, resembling *C. parvifolia* in the Chichijima Islands, differs in its reduced stellate hair density on both leaf surfaces and thinner leaves. *Callicarpa* × *chibusensis*, considered a natural hybrid between *C. subpubescens* and *C. boninensis*, is characterized by intermediate stellate hair densities—higher than in *C. subpubescens* but lower than in *C. boninensis*—on both leaf surfaces, smaller leaves than those of *C. subpubescens*, and a distinct flowering phenology relative to that of *C. boninensis*.

## Introduction

The genus *Callicarpa* L. (1753: 111), formerly classified under the family Verbenaceae (e.g., Chen & Gilbert 1994, Yamazaki 1993), is now assigned to the family Lamiaceae (e.g., Cantino *et al*. 1992, Li *et al*. 2016, Yonekura 2017). Comprising approximately 140 species of shrubs or trees—and rarely climbers—*Callicarpa* is primarily distributed in temperate and tropical regions (Yonekura 2017). The center of species diversity for *Callicarpa* lies in the Old World, particularly in China and Malesia, with approximately 50 species found in each region (Bramley 2013, Chen & Gilbert 1994). In Japan, 12 species, five varieties, and four natural hybrids have been reported (Yamamoto *et al*. 2023, Yamazaki 1993, Yonekura 2017). All Japanese species, except those in the Ogasawara Islands, including those in the Ryukyu Islands, are deciduous; only the species in the Ogasawara Islands are evergreen (Yonekura 2017). Furthermore, while other *Callicarpa* species worldwide are hermaphroditic, those found in the Ogasawara Islands are uniquely dioecious (Kawakubo 1998).

*Callicarpa* in the Ogasawara Islands represents a compelling example of adaptive radiation on oceanic islands (Ono 1991, Shimizu & Tabata 1991). Currently, three species are recognized: *Callicarpa subpubescens* Hook. et Arn. (1841: 305), *C. glabra* Koidz. (1918: en56), and *C. parvifolia* Hook. et Arn. (1841: 305). These species, found in the Chichijima Islands, inhabit a range of environments from dry scrubs to mesic forests. Among them, only *C. subpubescens* is distributed beyond the Chichijima Islands and occurs throughout the archipelago. Notably, *C. subpubescens* exhibits considerable morphological variation among populations in the Hahajima Islands (Kawakubo 1986). This species is a keystone component of the native vegetation in the Ogasawara Islands and serves as a focal species in revegetation efforts aimed at ecosystem restoration. However, *C. subpubescens* populations are declining due to the impacts of invasive alien species. Therefore, effective conservation planning requires a thorough understanding of the taxon, highlighting the need for a comprehensive taxonomic review of *Callicarpa* in the Ogasawara Islands.

To this end, we conducted genetic analyses, ecological surveys, and morphological measurements. Using 14 microsatellite markers (Mori *et al*. 2008, Sugai *et al*. 2019), we detected genetic differentiation within *C. subpubescens* populations in the Hahajima Islands, at a level comparable to that observed among the three *Callicarpa* species in the Chichijima Islands (Sugai *et al*. 2019). We additionally observed variation in flowering phenology within Hahajima Island (Sugai *et al*. 2019). Furthermore, by employing 17 microsatellite markers (Setsuko *et al*. 2018) and through a comprehensive field survey conducted across the Hahajima Islands, we identified four ecotypes and mapped their distributions (Setsuko *et al*. 2024a), along with their respective morphological traits and flowering phenologies (Setsuko *et al*. 2024a). In addition, we performed phylogenetic and population dynamics analyses of *Callicarpa* in the Ogasawara Islands, using RAD-seq to achieve higher resolution (Setsuko *et al*. 2024b). As a result, we identified one of the four ecotypes in the Hahajima Islands as the conventional species, *C. subpubescens*, whereas the remaining three were proposed as new taxa.

Herein, we describe these newly identified taxa as *Callicarpa boninensis* Sugai & Setsuko, *sp. nov*., *Callicarpa hahajimensis* Sugai & Setsuko, *sp. nov*., and *Callicarpa* × *chibusensis* Sugai & Setsuko, *hybr. nat. nov*., and provide detailed taxonomic information, including their distributions, habitats, etymologies, and preliminary IUCN conservation statuses. In addition, we present an identification key to the *Callicarpa* species in the Ogasawara Islands.

### Taxonomy

***Callicarpa boninensis*** Sugai & Setsuko *sp. nov*. Figure 1.

**Fig. 1.**
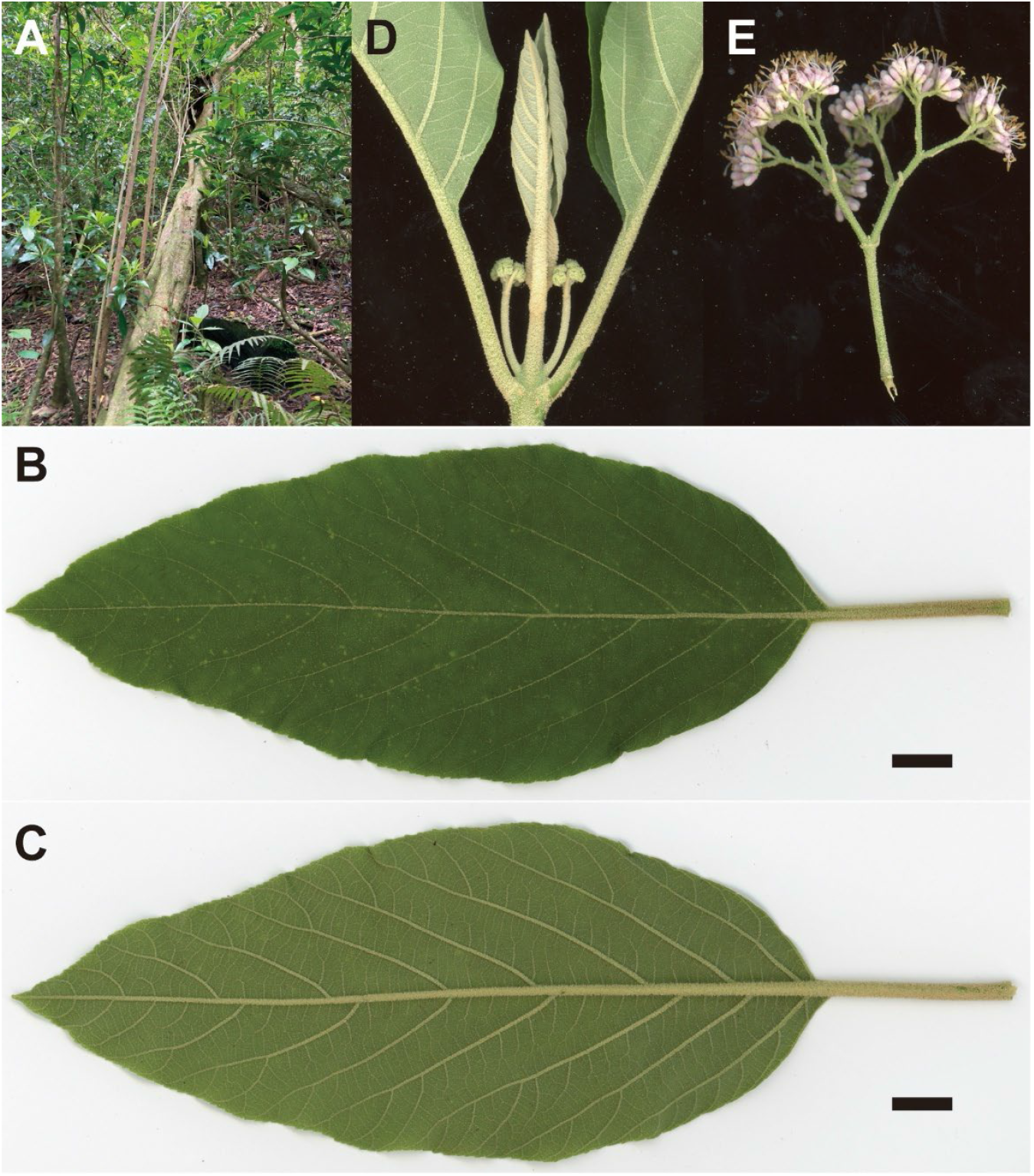
*Callicarpa boninensis* Sugai & Setsuko *sp. nov*. A, Habit. B, Adaxial leaf surface. C, Abaxial leaf surface. D, Branch apex with buds. E, Inflorescence (a female individual). Scale bars are 1 cm for B & C.

TYPE:—JAPAN. Tokyo Metropolis: Ogasawara Islands, Hahajima Island, Sekimon, 26°40’59.7”N, 142°09’43.0”E, elev. 248 m, 18 July 2023, *Kayo Hayama* (holotype: MAK472470!, isotypes: MAK472470! and TI00265245!).

### Diagnosis

*Callicarpa boninensis* is morphologically most similar to *C. subpubescens*, but can be distinguished by its persistent stellate hairs on mature leaves and longer petioles. The comparison among *C. subpubescens, C. boninensis, C. hahajimensis* and *C*. × *chibusensis* is shown in Table 1.

**Table 1.** Comparison of four *Callicarpa* taxa in the Hahajima Islands (Setsuko *et al*. 2024a).

| Characters | *C. subpubescens* | *C. boninensis* | *C. hahajimensis* | *C. × chibusensis* |
| --- | --- | --- | --- | --- |
| Tree height (m) | 1–5 | 3–11 | 0.3–2 | 2–5 |
| Blade thickness (mm) | 0.10 | 0.14 | 0.26 | 0.16 |
| Petiole length (cm) | 1–5 | 2–6 | 0.5–4 | 2–4 |
| Blade length (cm) | 7–21 | 7–16 | 2–11 | 8–13 |
| Blade width (cm) | 3–11 | 4–8 | 2–5 | 4–9 |
| Blade hairiness (upper) | glabrescent | stellate pilosa | sparsely stellate pilosa | glabrescent except sparsely stellate pubescent on midrib and lateral veins |
| Blade hairiness (lower) | glabrescent | densely stellate pubescent | stellate pubescent | sparsely stellate pubescent |
| Flowering time (range, peak) | Jun.–Jul., Jul. | Jul.–Dec., Oct. | Jul.–Jan., Aug. & Nov. | Jun.–Aug, Jul. |

Evergreen trees, typically 3–11 m tall, shorter on limestone substrates. Branches terete, with elliptic lenticels and raised leaf scars, densely covered with yellowish-brown soft stellate tomentose when young, later glabrescent. Leaves opposite, thickly chartaceous; petioles 2–6 cm long, stellate-pubescent; blades elliptic, oblong, or ovate, 7–16 cm long, 4–8 cm wide, apex acute or acuminate, base obtuse, shortly attenuated into petiole, margins serrulate on upper half or subentire, upper surface stellate-pilose, lower surface densely stellate-pubescent with a raised midrib and 7–8 pairs of obscure lateral veins. Flowers from July to December. Inflorescences axillary, dichasial cymes, densely many-flowered, 3–4.5 cm long, 2–3.5 cm wide, peduncle and rachis densely stellate tomentose. Pedicel ca. 1 mm long, glandular. Bracts lanceolate, apex obtuse, ca. 2 mm. Calyx cup-shaped, ca. 1.5 mm long, glandular, shallowly four-lobed, lobes widely deltate, obtuse, ca. 0.5 mm long. Corolla funnel-form, pale purple, ca. 5 mm long, sparsely glandular, four-lobed, lobes orbicular, ca. 2 mm long. Dioecious. Four stamens, exserted, ca. 7 mm long, anther ellipsoid, ca. 1 mm long, glandular-punctate, longitudinally dehiscent throughout. Style filiform, ca. 7 mm long (but 0.5–1.5 mm in males), stigma capitate. Fruit a drupe, globose, purple, ca. 3 mm in diameter.

### Additional specimens examined (paratype)

JAPAN. Tokyo Metropolis: Ogasawara Islands, **Hahajima Island**: Sekimon, 26°40’39.2”N, 142°09’26.8”E, 24 June 2006, *Keigo Mori* (MAK472402!); loc. cit., 26°40’39.5”N, 142°09’27.4”E, 4 October 2006, *K. Mori* (MAK472427!); loc. cit., 26°40’44.4”N, 142°09’31.7”E, 4 October 2006, *K. Mori* (MAK472428!, MAK472429!, MAK472430!, MAK472431! and MAK472432!); loc. cit., 26°40’44.9”N, 142°09’32.2”E, 4 October 2006, *K. Mori* (MAK472433!, MAK472434!, MAK472435!, MAK472436! and MAK472437!); loc. cit., 26°41’04.6”N, 142°09’38.6”E, 5 October 2006, *K. Mori* (MAK472412!); loc. cit., 26°41’05.9”N, 142°09’38.4”E, 5 October 2006, *K. Mori* (MAK472413!); loc. cit., 26°41’05.3”N, 142°09’39.8”E, 5 October 2006, *K. Mori* (MAK472414!); loc. cit., 26°41’05.0”N, 142°09’39.9”E, 5 October 2006, *K. Mori* (MAK472415!); loc. cit., 26°41’03.5”N, 142°09’39.8”E, 5 October 2006, *K. Mori* (MAK472416!); loc. cit., 26°41’03.2”N, 142°09’40.1”E, 5 October 2006, *K. Mori* (MAK472418!, MAK472419!, MAK472421!, MAK472422!, MAK472423! and MAK472424!); loc. cit., 26°41’02.9”N, 142°09’40.1”E, 5 October 2006, *K. Mori* (MAK472425!); loc. cit., 26°40’21.5”N, 142°09’21.6”E, 5 October 2006, *K. Mori* (MAK472446!); loc. cit.,26°40’25.1”N, 142°09’20.5”E, 5 October 2006, *K. Mori* (MAK472447! and MAK472458!); loc. cit., 26°40’20.1”N, 142°09’21.0”E, 18 June 2007, *K. Mori* (MAK472366!); loc. cit., 26°40’25.1”N, 142°09’20.5”E, 18 June 2007, *K. Mori* (MAK472367!); loc. cit., 26°40’52.5”N, 142°09’35.0”E, 18 June 2007, *K. Mori* (MAK464596! and MAK472467!); loc. cit., 17 March 1972, *Yasuichi Momiyama et al*. (MAK125161!); loc. cit., 26°40’24.9”N, 142°09’20.2”E, 29 June 2024, *Shota Sakaguchi et al*. (KYO00029254! and KYO00029255!); loc. cit., 26°40’45.5”N, 142°09’34.1”E, 29 June 2024, *S. Sakaguchi et al*. (KYO00029258! and KYO00029259!); Mt. Chibusa, 26°39’27.6”N, 142°09’45.9”E, 6 October 2006, *K. Mori* (MAK472275!); loc. cit., 26°39’23.8”N, 142°09’52.7”E, 6 October 2006, *K. Mori* (MAK472277!); loc. cit., 26°39’24.6”N, 142°09’51.7”E, 6 October 2006, *K. Mori* (MAK464585!); loc. cit., 26°39’23.8”N, 142°09’52.6”E, June 2014, *Satoshi Narita* and *Hidetoshi Kato* (MAK472496!). **Anejima Island**: 24 August 1980, *Mikio Ono et al*. (MAK184059!); loc. cit., 18 July 1992, *Takaya Yasui* (MAK268398!).

### Distribution

Japan. Tokyo Metropolis: Ogasawara Islands, Hahajima and Anejima Islands

### Habitat

Mesic forests in the Hahajima Islands dominated by *Elaeocarpus photiniifolia* Hook. et Arn., *Pisonia umbellifera* (J.R. et G.Forst.) Seem., *Ardisia sieboldii* Miq., and *Bischofia javanica* Blume.

### Etymology

The specific epithet *Boninensis* refers to its type locality: the Bonin (Ogasawara) Islands.

### Japanese name

Ogasawara-murasaki

### Conservation status

Genetically similar populations exist in the Mukojima and Volcano Islands (Setsuko *et al*. 2024a, Sugai *et al*. 2019); however, morphological characteristics and phenology remain insufficiently understood, preventing definitive species identification. Currently, this species is known only from Hahajima Island, which has an area of approximately 20 km^2^. The mesic forests on Hahajima Island have been invaded by the alien species, *B. javanica*, likely leading to a decline in the populations of *C. boninensis*. Consequently, we assign its conservation status as vulnerable (VU) following IUCN Red List criteria (IUCN 2012) under criteria D1, indicating the number of mature individuals is fewer than 1000, and D2, signifying the area of occupancy is less than 20 km^2^.

***Callicarpa hahajimensis*** Sugai & Setsuko *sp. nov*. Figure 2.

**Fig. 2.**
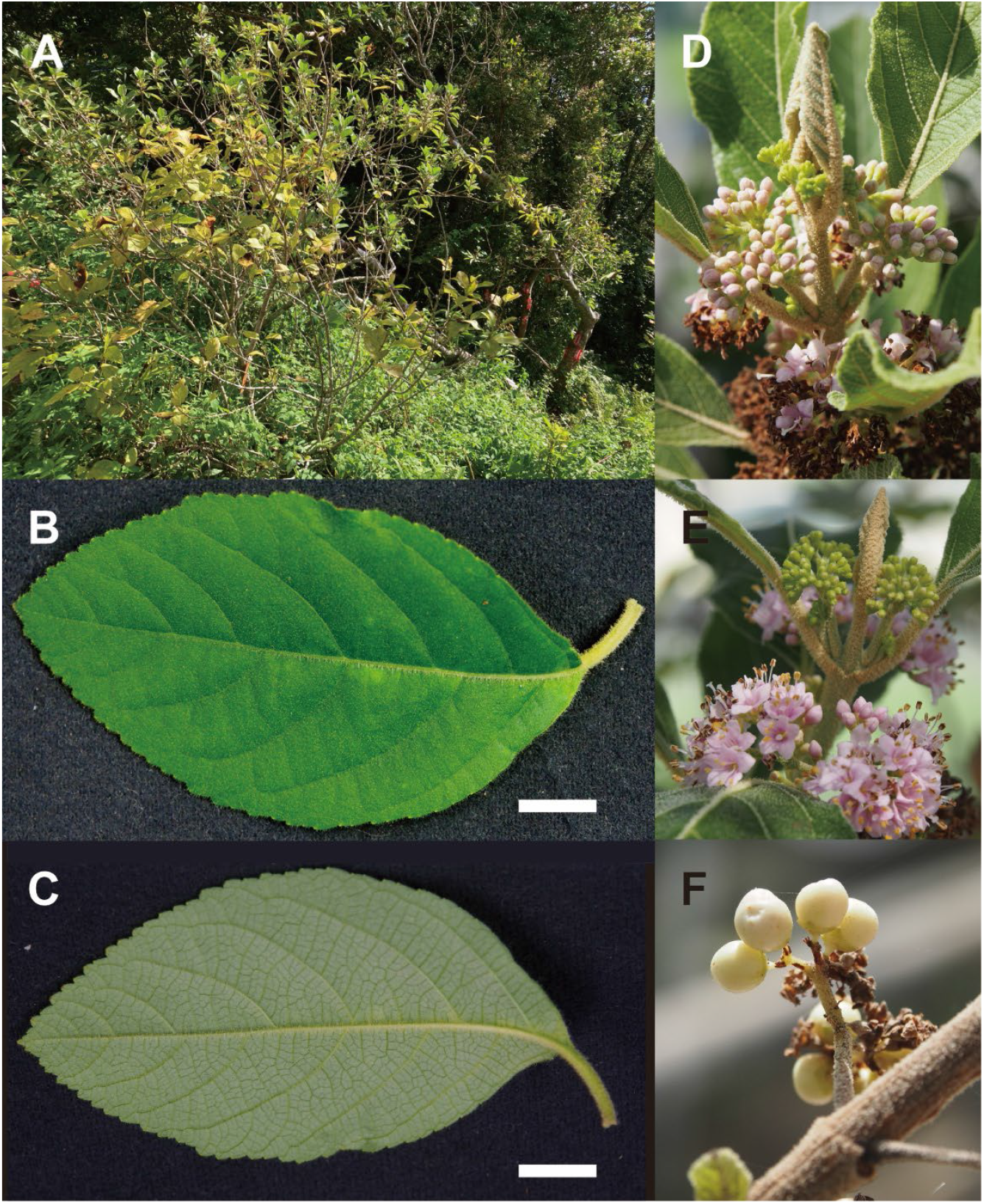
*Callicarpa hahajimensis* Sugai & Setsuko *sp. nov*. A, Habit. B, Adaxial leaf surface. C, Abaxial leaf surface. D, Branch apex with buds and flowers. E, Inflorescences (a female individual). F, Fruits. Scale bars are 1 cm for B & C.

TYPE:—JAPAN. Tokyo Metropolis: Ogasawara Islands, Hahajima Island, Mt. Kensaki, 26°38’47.5”N, 142°09’57.7”E, elev. 206 m, 18 July 2023, *K. Hayama* (holotype: MAK472471!, isotypes: MAK472471! and TI00265246!).

### Diagnosis

*Callicarpa hahajimensis* is morphologically most similar to *C. parvifolia* but can be distinguished by its lower density of stellate hairs on both leaf surfaces and thinner leaves. The comparison among *C. subpubescens, C. boninensis, C. hahajimensis* and *C*. × *chibusensis* is shown in Table 1.

Evergreen shrubs, 0.3–2 m tall, often caespitose. Branches terete, with elliptic lenticels and raised leaf scars, densely covered with yellowish-brown soft stellate tomentose when young, later glabrescent. Leaves opposite, thickly chartaceous; petioles 0.5–4 cm long, stellate-pubescent; blades elliptic, broadly elliptic, or ovate, 2–11 cm long, 2–5 cm wide, apex acute, base obtuse, shortly attenuated into petiole, margins serrulate on upper half, upper surface sparsely stellate-pilose, lower surface stellate-pubescent with a raised midrib and 5–8 pairs of obscure lateral veins. Flowers from July to January. Inflorescences axillary, dichasial cymes, densely many-flowered, 3–4 cm long, 2–3.5 cm wide, peduncle and rachis densely stellate tomentose. Pedicel ca. 1 mm long, glandular. Bracts lanceolate, apex obtuse, ca. 2 mm. Calyx cup-shaped, ca. 1.5 mm long, glandular, shallowly four-lobed, lobes widely deltate, obtuse, ca. 0.5 mm long. Corolla funnel-form, pale purple, ca. 5 mm long, sparsely glandular, four-lobed, lobes orbicular, ca. 2 mm long. Dioecious. Four stamens, exserted, ca. 7 mm long, anther ellipsoid, ca. 1 mm long, glandular-punctate, longitudinally dehiscent throughout. Style filiform, ca. 7 mm long (but ca. 0.5 mm in males), stigma capitate. Fruit a drupe, globose, purple or white, ca. 3 mm in diameter.

### Additional specimens examined (paratype)

JAPAN. Tokyo Metropolis: Ogasawara Islands, **Hahajima Island**: Mt. Kensaki, 26°38’43.4”N, 142°09’55.6”E, 11 November 2006, *H. Kato* (MAK472459! and MAK472460!); loc. cit., 26°38’45.1”N, 142°09’56.7”E, 11 November 2006, *H. Kato* (MAK4724619!); loc. cit., 26°38’45.8”N, 142°09’56.3”E, 11 November 2006, *H. Kato* (MAK472462! and MAK472463!); loc. cit., 26°38’45.9”N, 142°09’56.4”E, 11 November 2006, *H. Kato* (MAK472464!); loc. cit., 26°38’43.4”N, 142°09’55.6”E, 17 June 2007, *K. Mori* (MAK464593!); loc. cit., 26°38’44.4”N, 142°09’56.1”E, 17 June 2007, *K. Mori* (MAK464594!); loc. cit., 26°38’45.1”N, 142°09’56.7”E, 17 June 2007, *K. Mori* (MAK472317! and MAK472318!); loc. cit., 26°38’45.4”N, 142°09’56.8”E, 17 June 2007, *K. Mori* (MAK472319!); loc. cit., 26°38’45.5”N, 142°09’56.9”E, 17 June 2007, *K. Mori* (MAK472320! and MAK472321!); loc. cit., 26°38’45.8”N, 142°09’56.9”E, 17 June 2007, *K. Mori* (MAK472322! and MAK472323!); loc. cit., 26°38’44.5”N, 142°09’56.2”E, 17 June 2007, *K. Mori* (MAK472324! and MAK472325!); loc. cit., 26°38’44.5”N, 142°09’56.2”E, 17 June 2007, *K. Mori* (MAK472326!); loc. cit., 26°38’44.4”N, 142°09’56.0”E, 17 June 2007, *K. Mori* (MAK472327!); loc. cit., 26°38’44.5”N, 142°09’56.0”E, 17 June 2007, *K. Mori* (MAK472328!); loc. cit., 26°38’44.6”N, 142°09’55.9”E, 17 June 2007, *K. Mori* (MAK472329!, MAK472330! and MAK472331!); loc. cit., 26°38’44.5”N, 142°09’56.0”E, 17 June 2007, *K. Mori* (MAK472332!); loc. cit., 26°38’45.3”N, 142°09’56.9”E, June 2014, *S. Narita* and *H. Kato* (MAK472476!); loc. cit., 26°38’44.6”N, 142°09’56.2”E, June 2014, *S. Narita* and *H. Kato* (MAK472477!); loc. cit., 26°38’44.7”N, 142°09’56.0”E, June 2014, *S. Narita* and *H. Kato* (MAK472478!); Mt. Kensaki-Mt. Chibusa, 13 March 1988, *Motomi Ito et al*. (MAK299166!); Minamizaki, 26°37’23.3”N, 142°10’37.6”E, 15 September 2023, *K. Hayama* (MAK472474!); Mt. Higashi, 26°41’54.2”N, 142°08’53.1”E, 15 September 2023, *K. Hayama* (MAK472475!). **Imoutojima Island**: 26°33’35.6”N, 142°12’39.5”E, 23 June 2006, *K. Mori* and *H. Kato* (MAK464583!, MAK464584!, MAK472301!, MAK472302!, MAK472303!, MAK472304!, MAK472305!, MAK472306!, MAK472307! and MAK472308!); loc. cit., 26°33’35.2”N, 142°12’39.2”E, 15 June 2014, *Suzuki Setsuko* and *H. Kato* (MAK472480!); loc. cit., 26°33’35.7”N, 142°12’39.5”E, 15 June 2014, *S. Setsuko* and *H. Kato* (MAK472481!); loc. cit., 26°33’35.9”N, 142°12’39.5”E, 15 June 2014, *S. Setsuko* and *H. Kato* (MAK472482!); loc. cit., 26°33’35.3”N, 142°12’39.5”E, 15 June 2014, *S. Setsuko* and *H. Kato* (MAK472483!); loc. cit., 26°33’33.5”N, 142°12’34.5”E, 15 June 2014, *S. Setsuko* and *H. Kato* (MAK472484!); loc. cit., 26°33’36.0”N, 142°12’39.5”E, 15 June 2014, *S. Setsuko* and *H. Kato* (MAK472485!); loc. cit., 24 August 1980, *M. Ono et al*. (MAK183983!). **Meijima Island**: 24 August 1980, *M. Ono et al*. (MAK184005!).

### Distribution

Japan. Tokyo Metropolis: Ogasawara Islands, Hahajima, Imoutojima, and Meijima Islands

### Habitat

Dry scrubs in the Hahajima Islands, with forests dominated by *Rhaphiolepis indica* (L.) Lindl. var. *umbellata* (Thunb.) H.Ohashi, *Planchonella obovata* (R.Br.) Pierre, *Syzygium cleyerifolium* (Yatabe) Makino, and *Pandanus boninensis* Warb.

### Etymology

The specific epithet *hahajimensis* refers to its type locality: Hahajima Island.

### Japanese name

Hahajima-murasaki

### Conservation status

The dry scrubs in the Hahajima Islands have a limited local distribution. On Hahajima Island, these scrubs are under pressure from alien species, such as *Leucaena leucocephala* (Lam.) de Wit, *Kalanchoe pinnata* (Lam.) Pers. and *Bidens pilosa* L. var. *radiata* Sch. Bip., and have been invaded by *B. javanica*. In contrast, the satellite islands of Hahajima Island remain unaffected by alien species. Consequently, we assign its conservation status as endangered (EN) following IUCN Red List criteria (IUCN 2012) under criterion D, indicating that the number of mature individuals is fewer than 250.

***Callicarpa* × *chibusensis*** Sugai & Setsuko *hybr. nat. nov*. Figure 3.

**Fig. 3.**
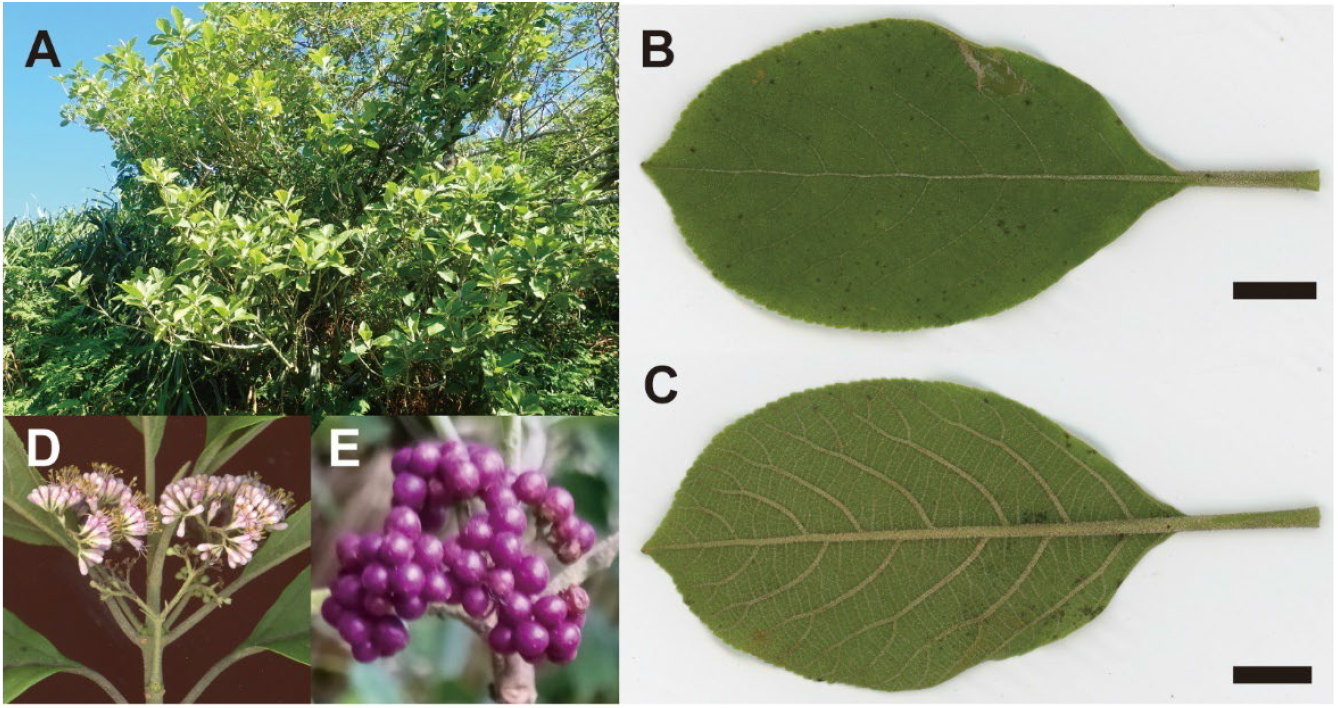
*Callicarpa* × *chibusensis* Sugai & Setsuko *hybr. nat. nov*. A, Habit. B, Adaxial leaf surface. C, Abaxial leaf surface. D, Inflorescences (a female individual). E, Fruits. Scale bars are 1 cm for B & C.

TYPE:—JAPAN. Tokyo Metropolis: Ogasawara Islands, Hahajima Island, Mt. Chibusa, 26°39’29.1”N, 142°09’44.5”E, elev. 413 m, 30 April 2023, *S. Setsuko* and *K. Hayama* (holotype: MAK472472!, isotypes: MAK472472! and TI00265244!).

### Diagnosis

*Callicarpa* × *chibusensis* is morphologically similar to both *C. subpubescens* and *C. boninensis*, but can be distinguished by its higher and lower density of stellate hairs on both leaf surfaces compared to *C. subpubescens* and *C. boninensis*, respectively, as well as smaller leaves than those of *C. subpubescens*. It also exhibits a distinct flowering phenology compared to that of *C. boninensis*. This plant is considered a natural hybrid between *C. subpubescens* and *C. boninensis*. The comparison among *C. subpubescens, C. boninensis, C. hahajimensis* and *C*. × *chibusensis* is shown in Table 1.

Evergreen shrubs, 2–5 m tall. Branches terete, with elliptic lenticels and raised leaf scars, densely covered with yellowish-brown soft stellate tomentose when young, later glabrescent. Leaves opposite, thickly chartaceous; petioles 2–4 cm long, stellate-pubescent; blades elliptic, oblong, broadly elliptic, or ovate, 8–13 cm long, 4–9 cm wide, apex acute or acuminate, base obtuse, shortly attenuated into petiole, margins serrulate on upper half, upper surface glabrescent except sparsely stellate-pubescent on midrib and lateral veins, lower surface sparsely stellate-pubescent with a raised midrib and 5–7 pairs of obscure lateral veins. Flowers from June to August. Inflorescences axillary, dichasial cymes, densely many-flowered, 3–5.5 cm long, 2.5–4 cm wide, peduncle and rachis densely stellate tomentose. Pedicel ca. 1 mm long, glandular. Bracts lanceolate, apex obtuse, ca. 2 mm. Calyx cup-shaped, ca. 1.5 mm long, glandular, shallowly four-lobed, lobes widely deltate, obtuse, ca. 0.5 mm long. Corolla funnel-form, pale purple, ca. 5 mm long, sparsely glandular, four-lobed, lobes orbicular, ca. 2 mm long. Dioecious. Four stamens, exserted, ca. 7 mm long, anther ellipsoid, ca. 1 mm long, glandular-punctate, longitudinally dehiscent throughout. Style filiform, ca. 7 mm long (but 0.5–1 mm in males), stigma capitate. Fruit a drupe, globose, purple, ca. 3 mm in diameter.

### Additional specimens examined (paratype)

JAPAN. Tokyo Metropolis: Ogasawara Islands, **Hahajima Island**: Mt. Chibusa, 26°39’33.4”N, 142°09’39.9”E, 22 June 2006, *K. Mori* (MAK472398!); loc. cit., 26°39’31.8”N, 142°09’42.2”E, 6 October 2006, *K. Mori* (MAK464586!); loc. cit., 26°39’32.2”N, 142°09’37.5”E, 6 October 2006, *K. Mori* (MAK472257!); loc. cit., 26°39’33.4”N, 142°09’39.9”E, 6 October 2006, *K. Mori* (MAK472258!); loc. cit., 26°39’31.7”N, 142°09’42.5”E, 6 October 2006, *K. Mori* (MAK472259!); loc. cit., 26°39’31.4”N, 142°09’42.4”E, 6 October 2006, *K. Mori* (MAK472260!); loc. cit., 26°39’27.9”N, 142°09’46.0”E, 6 October 2006, *K. Mori* (MAK472261!); loc. cit., 26°39’27.2”N, 142°09’46.0”E, 6 October 2006, *K. Mori* (MAK472262!); loc. cit., 26°39’25.1”N, 142°09’51.4”E, 6 October 2006, *K. Mori* (MAK472263!); loc. cit., 26°39’24.1”N, 142°09’52.1”E, 6 October 2006, *K. Mori* (MAK472264!); loc. cit., 26°39’22.4”N, 142°09’53.8”E, 6 October 2006, *K. Mori* (MAK472265!); loc. cit., 26°39’33.1”N, 142°09’38.6”E, 6 October 2006, *K. Mori* (MAK472266!); loc. cit., 26°39’33.1”N, 142°09’39.0”E, 6 October 2006, *K. Mori* (MAK472267! and MAK472268!); loc. cit., 26°39’33.5”N, 142°09’39.4”E, 6 October 2006, *K. Mori* (MAK472269!); loc. cit., 26°39’31.7”N, 142°09’42.5”E, 6 October 2006, *K. Mori* (MAK472270!); loc. cit., 26°39’31.4”N, 142°09’42.4”E, 6 October 2006, *K. Mori* (MAK472271!); loc. cit., 26°39’29.1”N, 142°09’44.0”E, 6 October 2006, *K*. Mori (MAK472272!); loc. cit., 26°39’25.7”N, 142°09’49.9”E, 6 October 2006, *K. Mori* (MAK472276!); loc. cit., 19 November 1970, *M. Ono* and *Sumiko Kobayashi* (MAK123833!); loc. cit., 29–30 June 1975, *S. Kobayashi* (MAK138410! and MAK138409!); loc. cit., 19 June 1979, *S. Kobayashi* and *Yuji Ohmori* (MAK180608! and MAK180609!); loc. cit., 19 November 1997, *Jin Murata et al*. (MAK293812!); loc. cit., 13 December 1998, *Tetsuo Ohi* (MAK306701!); loc. cit., 28 April 2001, *H. Kato* (MAK319233!); loc. cit., 26°39’21.6”N, 142°09’50.4”E, 30 June 2024, *S. Sakaguchi et al*. (KYO00029260!, KYO00029261!, KYO00029262!, KYO00029263! and KYO00029264!); Mt. Chibusa-Sakaigatake, 25 March 1971, *M. Ono* and *S. Kobayashi* (MAK124956! and MAK124957!).

### Distribution

Japan. Tokyo Metropolis: Ogasawara Islands, Hahajima Island **Habitat**:—Mesic scrubs and the edge of mesic forests (cloud forests) on the main ridge at >350 m elevation on Hahajima Island. The forests are dominated by *Freycinetia formosana* Hemsl. var. *boninensis* Nakai, *Dendrocacalia crepidifolia* (Nakai) Nakai, *Fatsia oligocarpella* Koidz., and *Melastoma tetramerum* Hayata var. *pentapetalum* Toyoda.

### Etymology

The specific epithet *chibusensis* refers to its type locality: Mt. Chibusa.

### Japanese name

Chibusa-shima-murasaki

### Conservation status

As a natural hybrid, this species inhabits slightly different environments compared to its parent species and is exclusively found in the cloud forests at higher elevations on Hahajima Island. Although the population is not small, it is likely to be declining due to the encroachment of the alien species *B. javanica*. Therefore, we assign its conservation status as vulnerable (VU) following IUCN Red List criteria (IUCN 2012) under criteria D1, indicating the number of mature individuals is fewer than 1000, and D2, signifying the area of occupancy is less than 20 km^2^.

**Key for *Callicarpa* species in the Ogasawara Islands** [partly based on Yamazaki (1993) and Yonekura (2017)]

1a Deciduous shrubs. Hermaphrodite….. Species in Japan, except the Ogasawara Islands
1b Evergreen shrubs or trees. Dioecy 2
2a Branches and leaves glabrous *C. glabra*
2b Young branches and petioles densely stellate-hairy 3
3a Leaves thickly chartaceous, lower surface densely stellate tomentose. Shrubs ca. 1 m tall. Leaf Flowers June to August *C. parvifolia*
3b Leaves chartaceous 4
4a Leaf surfaces sparsely pubescent or glabrous. Shrubs 1–5 m tall. Flowers May to August *C. subpubescens*
4b Leaf lower surface stellate-pubescent 5
5a Shrubs 0.3–2.0 m tall, often caespitose. Flowers over a prolonged period from July to January *C. hahajimensis*
5b Flowers summer or autumn 6
6a Shrubs 2–5 m tall. Flowers June to August *C*. × *chibusensis*
6b Trees 3–11 m tall, often taller on well-developed soils. Flowers July to December *C. boninensis*

## Acknowledgements

The authors are grateful to the Metropolis of Tokyo, the Ministry of the Environmental Government of Japan, and the Forestry Agency of Japan for allowing this study. The authors thank Dr. Shota Sakaguchi for providing photos for this study. This work was funded by Grant-in-Aid for Science Research from the Japanese Society for Promotion of Science (JP26290073, JP15K07203, JP21K05694 and JP21KK0131), the Environment Research and Technology Development Fund of the Ministry of the Environment, Japan (4-1402), and “Shimane University Grants for Joint Research Project led by Female Researchers” under the MEXT “Initiative for Realizing Diversity in the Research Environment (Collaboration Type).”

